# Towards high-throughput parallel imaging and single-cell transcriptomics of microbial eukaryotic plankton

**DOI:** 10.1101/2023.03.29.534285

**Authors:** Vesna Grujcic, Sami Saarenpää, John Sundh, Bengt Sennblad, Benjamin Norgren, Meike Latz, Stefania Giacomello, Rachel A Foster, Anders F Andersson

**Affiliations:** Department of Gene Technology, School of Engineering Sciences in Chemistry, Biotechnology and Health,SciLifeLab, KTH Royal Institute of Technology, Stockholm, Sweden; Department of Biochemistry and Biophysics, National Bioinformatics Infrastructure Sweden, SciLifeLab, Stockholm University, Box 1031, SE-17121 Solna, Sweden; Dept of Cell and Molecular Biology, National Bioinformatics Infrastructure Sweden, Science for Life Laboratory, Uppsala University, Husargatan 3, SE-752 37 Uppsala, Sweden; Department of Ecology, Environment and Plant Sciences, Stockholm University, Stockholm, 106 91 Sweden

**Author notes:** Corresponding authors: Anders F. Andersson and Vesna Grujcic.

**Keywords:** single-cell transcriptomics, MASC-seq, microbial eukaryotes, protists, plankton

## Abstract

Single-cell transcriptomics has the potential to provide novel insights into poorly studied microbial eukaryotes. Although several such technologies are available and benchmarked on mammalian cells, few have been tested on protists. Here, we optimized a microarray single-cell sequencing (MASC-seq) technology that generates microscope images of cells in parallel with capturing their transcriptomes. We tested the method on three species representing important plankton groups with different cell structures, the ciliate *Tetrahymena thermophila*, the diatom *Phaeodactylum tricornutum* and the dinoflagellate *Heterocapsa* sp.. Both the cell fixation and permeabilization steps were adjusted. For the ciliate and dinoflagellate, the number of transcripts of microarray spots with single cells were significantly higher than for background spots, and the overall expression patterns were correlated with that of bulk RNA, while for the much smaller diatom cells, it was not possible to separate single-cell transcripts from background. The MASC-seq method holds promise for investigating “microbial dark matter”, although further optimizations are necessary to increase the signal-to-noise ratio.

## Introduction

Planktonic microbial eukaryotes (protists) function as primary producers, grazers and parasites, and are important in carbon and nutrient cycling^1–4^. Currently, however, we know relatively little about the biology of most microbial eukaryotes and even less about how they interact and function at the ecosystem level^5^. The genome is the blueprint for understanding the physiology, ecology and evolution of an organism, but the majority of protists are difficult to culture and therefore obtaining genomic references for them has been challenging, and to date there are only a few reference genomes available^6, 7^. Shotgun metagenomics and binning allows recovering genomes without cultivation, and is effective for prokaryotes^8, 9^. However, due to the typically low relative abundance and higher complexity of eukaryotic genomes, this approach has so far had more limited success for protists (but see e.g.^10^).

Genomes of uncultivated plankton can also be obtained through single-cell sequencing^11^. Since eukaryotic genomes are complex, an attractive alternative is to sequence the transcriptome (RNA sequencing; RNA-seq), which allows characterization of expressed genes without having to sequence the sometimes vast non-coding regions that are common in eukaryotic genomes. Single-cell RNA sequencing (scRNA-seq) emerged about a decade ago, as a powerful tool in biomedical research to study cell-to-cell variability^12^ and has mainly been applied to mammalian cells. Applications of the method to cell types such as plants^13^ and other eukaryotes^14, 15^ are slowly emerging but still scarce, since the process requires vigorous adaptation of the protocol. Application of scRNA-seq to microbial eukaryotes has great potential for revealing ecophysiological properties from basic traits such as phototrophy, phagotrophy or osmotrophy, to more elaborate functions such as nutrient uptake, storage systems and interactions with other microbes^16, 17^.

To date, only a few different scRNA-seq protocols and methods have been tested on microbial eukaryotes, mostly due to difficulty of adapting protocols to extensive variation in cell size and cell surface structure. Recent studies using versions of SMART-seq showed performances comparable to bulk RNA-seq on bigger ciliate cells (50 - 100 μm) without hard cell walls^18, 19^; however, achieved only low gene recovery and high stochasticity of gene expression on small cells with cell walls (a haptophyte - 8 μm and a dinoflagellate - 15 μm)^20^. Another technology, the 10X Genomics Chromium platform widely used in biomedical research, showed results comparable to bulk RNA-seq on the photosynthetic unicellular algae *Chlamydomonas* with no cell wall^21^, indicating that cell size and cell wall are determining factors in application of scRNA-seq protocols to microbial eukaryotes.

In this study we applied MASC-seq (massive and parallel microarray sequencing)^22^, a scRNA-seq method that enables sequencing of hundreds of cells at the same time, and that links microscope images with the single cell transcriptome data. For this method, fixed cells are spread over a glass slide that contains array with 100 μm-sized spots of DNA capture probes with spot-specific indices. The cells are first imaged using a scanning microscope and then permeabilized, releasing their RNA out of the cells which binds to the probes on the array. cDNA is synthesized, harvested and sequenced, and, using the spot-specific barcode-sequences, cDNA sequences stemming from a specific spot (i.e., cell) can be linked to the microscope image of the corresponding cell. However, until now, the MASC-seq method has only been applied to mammalian cells.

The aim of this study was to test and adapt the MASC-seq method for application on unicellular eukaryotic plankton. We applied and optimized the method on three cultured plankton representatives, abundant in communities of aquatic environments, *Phaeodactylum tricornutum* (a diatom, silica and polysaccharide cell walls^23^), *Heterocapsa* sp. (a dinoflagellate, cellulose thecal plates^24^) and *Tetrahymena thermophila* (a ciliate, lipid membrane^25^) which all have different size and diverse cell surface structures common to plankton. We optimized several steps in the protocol to make it more suitable for planktonic cells and compared the results from MASC-seq generated single cell transcriptomes to bulk RNA sequencing.

## Results

### Adapting the MASC-seq method for eukaryotic single-celled plankton

In order to adapt the MASC-seq protocol to eukaryotic plankton, we focused on three species, each representing an important group of plankton: *P. tricornutum* (a diatom), *Heterocapsa sp.* (a dinoflagellate) and *T. thermophila* (a ciliate). The three species were cultured individually and cells were harvested at the exponential growth phase. The following steps of the MASC-seq protocol were optimized using these cells: cell fixation, cell permeabilization and cell attachment to the specialized microarray slide. These tests were performed on optimization slides and library preparation slides (Figure 1E-C) and based on the results we recommend the following to the existing protocol by Vickovic et al.^22^

**Figure 1.**
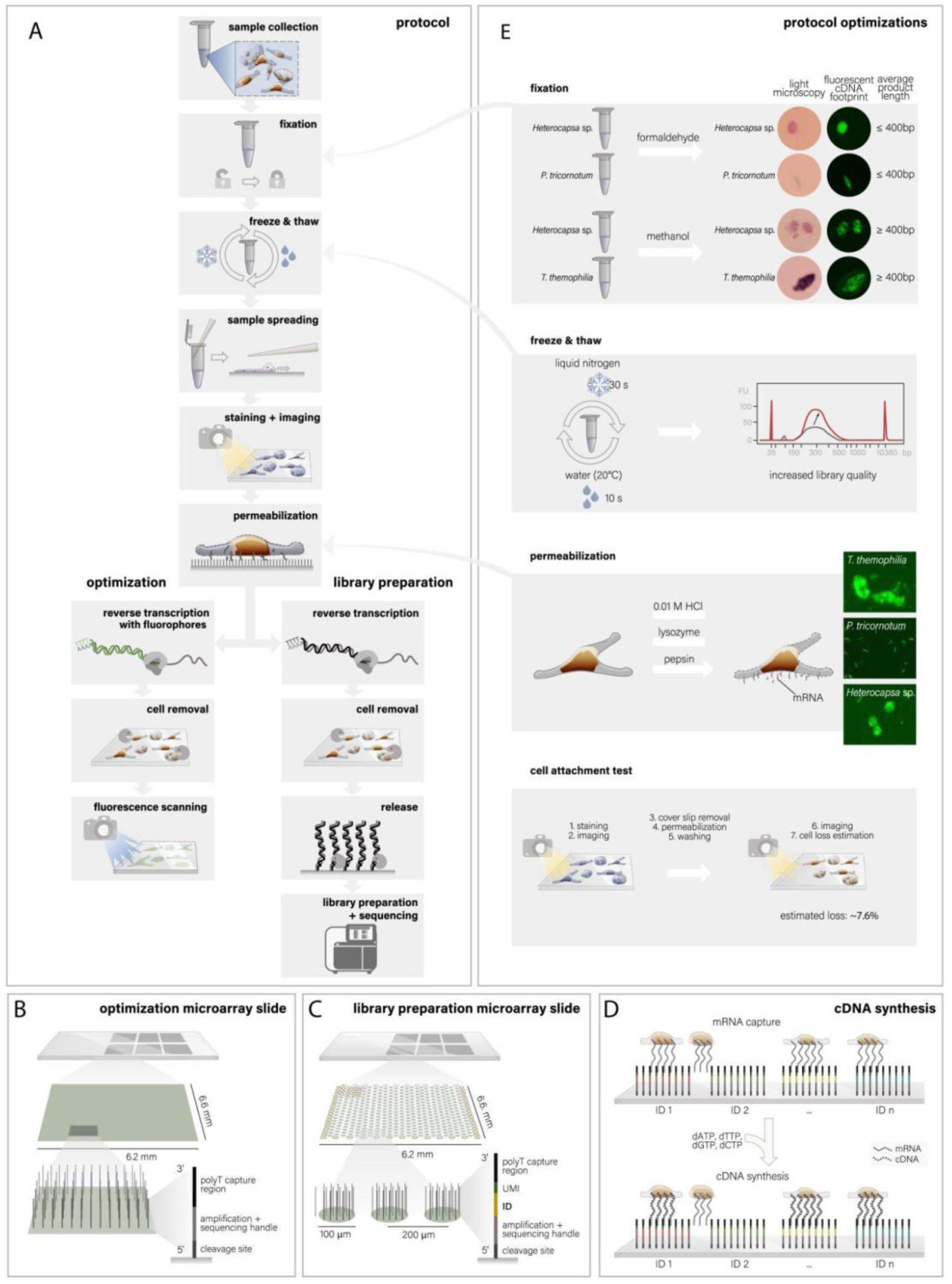
The different steps of the MASC-SEQ method. **A**) Samples are collected as described in the Methods section and fixed. Eukaryotic plankton that have hard cell wall covering undergo freeze-thaw cycling in liquid nitrogen to pre-permeabilize cell walls. Subsequent steps are attachment to the slide, staining, imaging and permeabilization. After permeabilization, polyadenylated transcripts leaking out of the cells are captured by the surface probes, which function as primers in the following reverse transcription reaction. Cells are removed, cDNA is enzymatically cleaved off and harvested, amplified and processed for Illumina sequencing (path to the right in panel A). **C**) The slides have six separate arrays that can accommodate different experiments. Each array contains ∼2000 spots with oligo-dT probes that have spot-specific barcodes (IDs) and UMIs. **D**) During cDNA synthesis, a cell positioned on top of a certain spot will get the spot-specific ID incorporated into its cDNA that can later be linked to the image of the spot. For optimization purposes, a special type of optimization arrays with oligo-dT probes covering the whole surface (**B**) can be used. Here, the reverse transcription is conducted using fluorescently labeled nucleotides, and after cell removal, the slide is scanned and the fluorescence cDNA footprint will correspond to the amount of cDNA synthesized (path to the left in panel A). Library preparation and sequencing are not conducted in this case. **E**) Optimizations conducted to make the MASC-seq protocol more suitable for single-celled eukaryotic plankton. From top to bottom: tested fixatives, formaldehyde (2%) and methanol (absolute) both gave positive results on optimization array slide, methanol produced longer final library molecules. Two different enzymes (pepsin and lysozyme) and acid (0.01 M HCL) were tested for cell permeabilization and only pepsin gave positive results. Freeze-thaw cycling in liquid nitrogen and room temperature (3 cycles) improved the amount of product in the final libraries. A cell attachment test was performed to estimate cell loss during the different steps of the protocol. Figure illustrations in A, B and C are adapted with permission from^36^, Springer Nature Limited.

#### Cell fixation

We tested how two different cell fixatives (formaldehyde and methanol) affect cDNA synthesis and final library quality. Here, we used cells of *P. tricornotum* and *Heterocapsa sp.* as models. The two methods showed comparable fluorescent cDNA signals on optimization slides, but, as expected, the Bioanalyzer traces of final sequencing libraries showed that methanol gave longer final library products (Figure 1E, Supplementary Figure 1). Consequently, we used methanol as the fixative of choice when preparing the libraries for sequencing. Related to the fixation step, we additionally compared prefixing the cells during the collection (using methanol as a fixative) with keeping them unfixed and then, during the first stages of the protocol, fixing them on the slide (as done by Vickovic et al.^22^). Our results indicated that cell morphology was improved when cells were prefixed during the collection, since the staining (see below) was more even and cells retained their shape better (Figure 1E, Supplementary figure 2).

#### Cell permeabilization

Vickovic et al.^22^ used pepsin for 30s when permeabilizing cells of the human adenocarcinoma cell line. In addition to pepsin, we tested lysozyme and 0.01 M HCL (Supplementary figure 1) which are both commonly used in catalyzed reporter deposition fluorescent *in situ* hybridization (CARD-FISH) assays for permeabilizing eukaryotic plankton^26^. However, of the three tested permeabilization methods, only pepsin rendered positive fluorescent cDNA signals on the optimization slides (**Figure 1E**). Cells of *P. tricornutum* and *Heterocapsa sp.* were permeabilized with pepsin for 5 min and *T. thermophila* for 1 min due to the difference in cell structures. *T. thermophila* has phospholipid membrane, similar to mammalian cells meaning shorter permeabilization time compared to *P. tricornutum* and *Heterocapsa sp.,* which both have hard cell walls.

We further compared solely using pepsin or combining it with freeze-thaw cycles in liquid nitrogen and room temperature water (details in Methods section). Freeze-thaw cycles increased concentrations of final libraries (Figure 1E, Supplementary figure 3) for *P. tricornutum* and *Heterocapsa sp.*, but resulted in disintegration of the less robust *T. thermophila* cells(Supplementary figure 4).

#### Cell attachment

After the cells have been imaged, they undergo different incubations and washing steps (see Methods section), which may lead to cell loss before their mRNA has been captured by the surface probes. To estimate the extent of cell loss, we used *Heterocapsa sp.* as a model and calculated that around 7% of the cells fall off during the different stages of the protocol (Figure 1E). This is an important aspect to consider when comparing hematoxylin and eosin (H&E) images and images of fluorescent cDNA synthesis.

### MASC-seq applied on individual plankton species

To evaluate quality and quantity of data produced with the MASC-seq method we sequenced four libraries, two for *T. thermophila* (Tet1 and Tet2), one for *Heterocapsa* sp. (Het) and one for *P. tricornotum* (Pha), using the optimized protocol. By manual inspection of the high-resolution microscope images of the arrays, we subdivided spots on each array into the following four groups: single cells (1 cell per spot), pairs of cells (2 cells), clusters of cells (>2 cells), and background (0 cells) (Figure 2). There was a general pattern that spots with more cells displayed higher expression (more expressed genes [i.e., expressed reference transcripts, see Methods section] and more unique transcripts) than those with fewer cells (Figure 2). Background spots had significantly lower expression than single cell spots in all libraries except *P. tricornotum*, and single-cell spots had significantly lower expression than double-cell spots in all libraries except *Heterocapsa* sp. (Figure 2).

**Figure 2.**
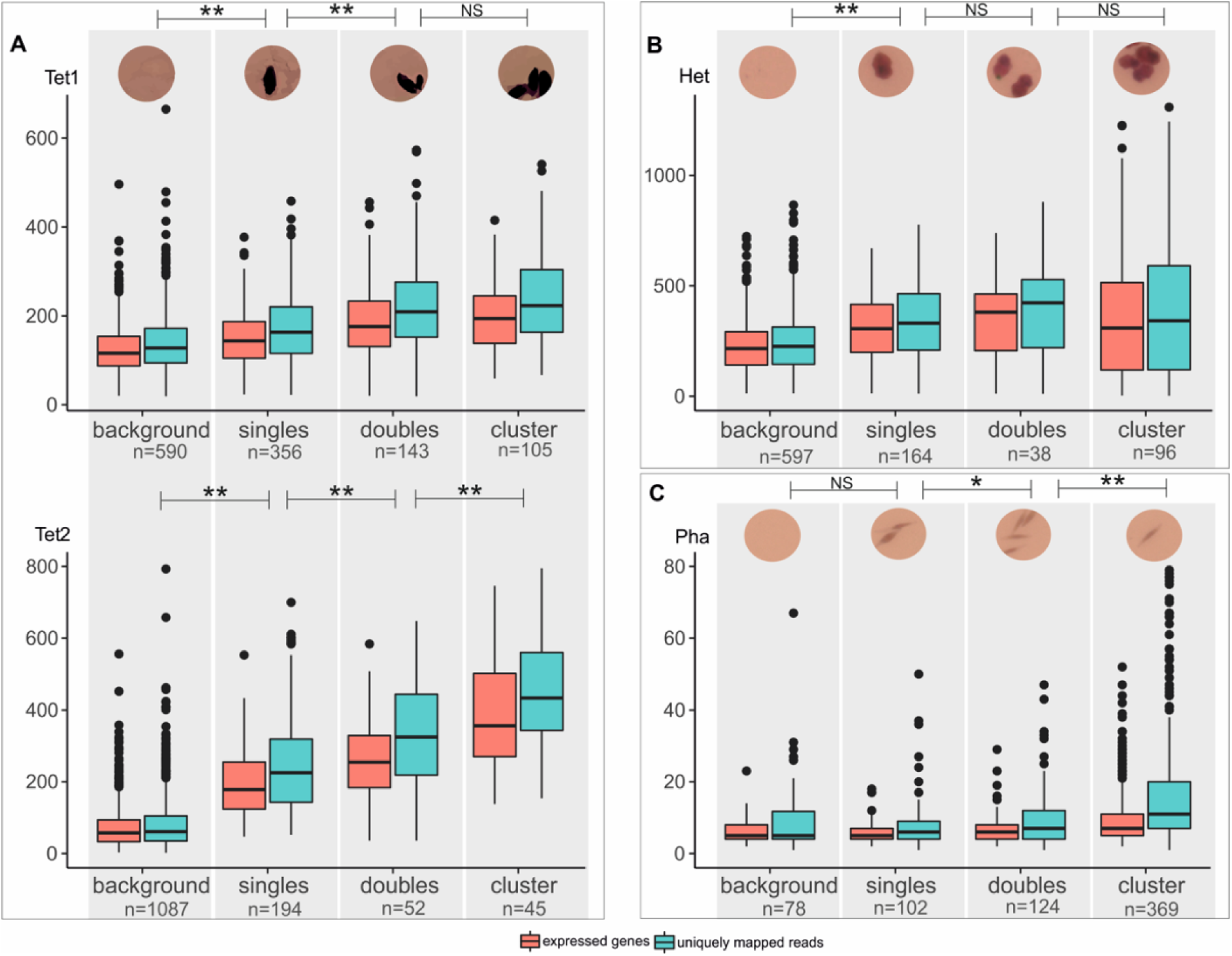
Number of expressed genes and unique transcripts per spot for different spot categories (background [empty spots], singles [one cell], doubles [two cells] and clusters [>2 cells]) identified based on the high-resolution images and MASC-seq sequencing libraries of **A**) *T. thermophila* (two libraries; Tet1 and Tet2), **B**) *Heterocapsa* sp. (Het) and **C**) *P. tricornutum* (Pha). An example image of a spot is shown above each spot category and species. The number of spots (n) for each category and library are given. Significance levels of Wilcoxon rank-sum tests comparing number of unique transcripts per spot between pairs of spot categories are indicated, with * = p-value < 0.05; ** = p-value < 0.0001; and NS = non-significant.

We compared the gene expression obtained from the cells on the array with that of total RNA extracted from the bulk cultures and pipetted onto the array (Figure 3). Samples for RNA were collected at the same time as cells for MASC-seq, and the amount of RNA was adjusted to correspond to the expected total amount of RNA in the ca. 2000 cells in the MASC-seq experiments (see below). The total expression (number of transcripts) per gene was significantly correlated between cells and RNA for all three species, but the linear correlations were stronger for the two *T. thermophila* replicates and *Heterocapsa* sp. (Pearson *r* = 0.64 - 0.73) than for *P. tricornotum* (*r* = 0.43), likely due to the much larger number of transcripts detected in the former. The correlations were also lower than between the two replicated MASC-seq experiments in *T. thermophila* (*r* = 0.88) (Figure 3), which may indicate that the cell permeabilization, cDNA synthesis and library preparation in this protocol probably give rise to some transcript-specific biases not present in the more complete lysis of cells in the bulk RNA preparations.

**Figure 3.**
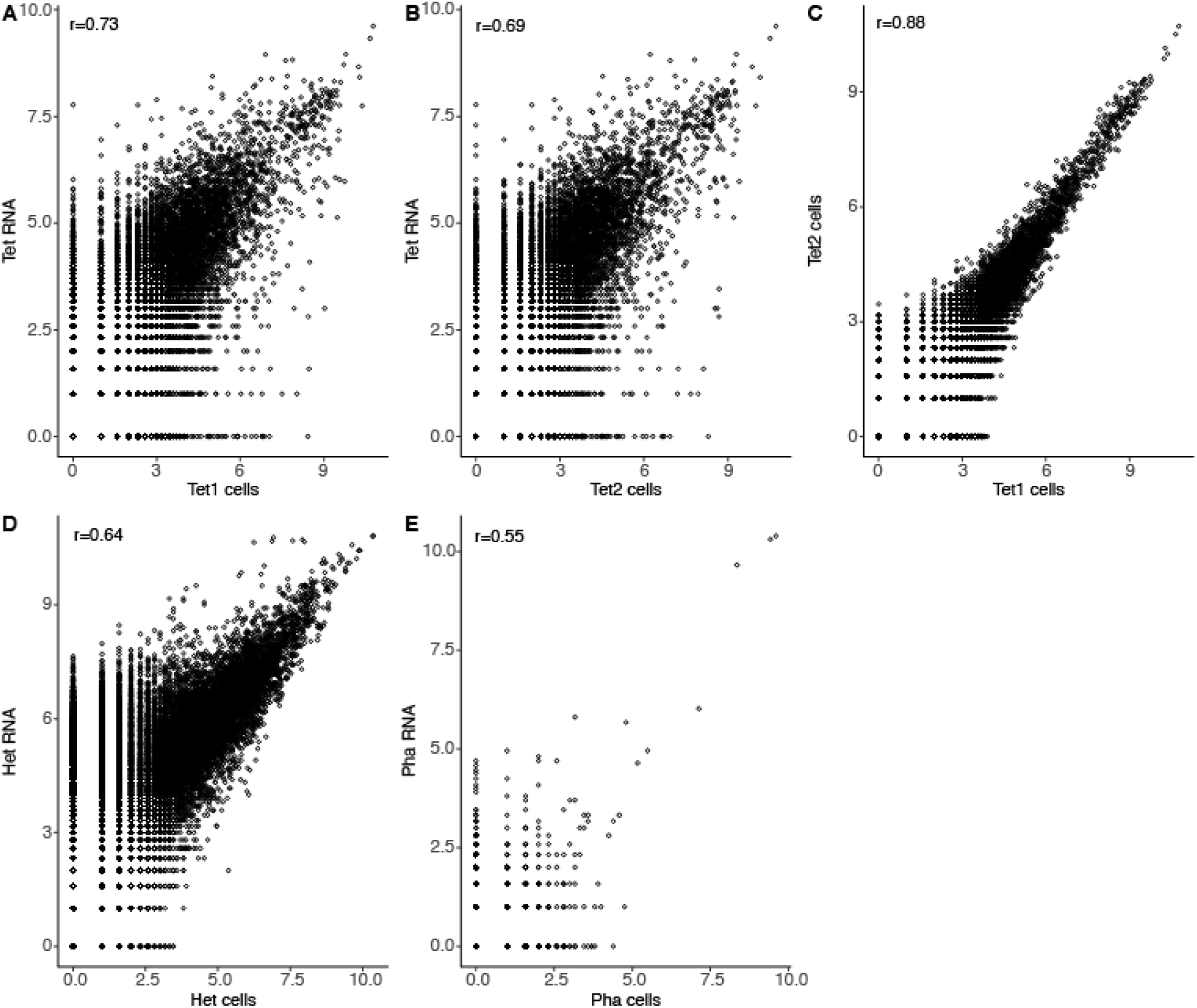
Scatter plots illustrating correlations between sequencing libraries. Each circle represents one gene, and its position indicates its total counts (number of unique transcripts) in the two libraries compared. **A-C)** *T. thermophila.* Bulk RNA added to the array vs. MASC-seq data (cells) for Tet1 (A) and Tet2 (B). MASC-seq data for Tet1 vs Tet2 (C). **D)** Bulk RNA added to the array vs. MASC-seq data for *Heterocapsa* sp. **E)** Bulk RNA added to the array vs. MASC-seq data for *P. tricornutum*. Pearson correlation coefficients (r) are given for each subplot.

The much lower number of expressed genes and transcripts per cell in *P. tricornotum* (on average 4.9 genes and 8.0 transcripts for single-cell spots) compared to *Heterocapsa* sp. (305 genes and 338 transcripts) and *T. thermophila* (148 and 194 genes, and 171 and 254 transcripts, for the two libraries) may partly be explained by the smaller size of *P. tricornotum* and the expected lower number of mRNA molecules inside each cell. *P. tricornotum* cells have a volume of 60 - 170 µm^3 27^ as compared to 1,000 - 4,000 µm^3^ and 15,000 µm^3^ for *Heterocapsa* sp.^27^ and *T. thermophila*^28^, respectively. In line with this, based on the amount of RNA that we could harvest from cultures of each cell type, we estimated that *P. tricornotum* had 1,250 – 6,250 mRNA molecules per cell compared to *Heterocapsa* sp. with 27,500 - 137,500 and *T. thermophila* with 46,490 - 232,450 molecules^20^ (Table 1). However, cell size cannot be the only explanation behind the variation in detected transcripts per cell, as illustrated by the variation between the two *T. thermophila* libraries. Other factors such as optimal cell fixation and cell permeabilization conditions, optimal amplification of library molecules, and sequencing depth are likely important, although much larger variability in scRNA transcripts was observed in protist cells compared to mammalian cells^20^.

**Table 1.** Cell sizes and estimated number of total RNA and mRNA molecules per cell. The number of RNA molecules calculated from the amount of total RNA extracted and the number of cells used for the RNA extractions. ^a^Assuming RNA extraction efficiency was 50% and an average mRNA transcript length of 1000 nt, ^b^Assuming 1–5% of total RNA was mRNA. (according to Liu et al. 2017)

| | Cell biovolume<br>($\mu\text{m}^3$) | Total RNA molecules<br>per cell<sup>a</sup> | mRNA molecules per<br>cell<sup>b</sup> |
| --- | --- | --- | --- |
| *P. tricornotum* | 60 - 170 $\mu\text{m}^3$ | 125 000 | 1250 - 6290 |
| *Heterocapsa sp.* | 1000 - 4000 $\mu\text{m}^3$ | 2 750 000 | 27 500 - 137 500 |
| *T. thermophila* | 15000 $\mu\text{m}^3$ | 4 649 000 | 46 490 - 232 450 |

In order to test the effect of sequencing depth on the number of detected transcripts per cell, we subsampled the datasets to different levels (Figure 4). This showed that a plateau in the detected number of transcripts and genes was reached before the full library had been analyzed for *P. tricornutum* (Figure 4E-F) while for *Heterocapsa* sp., and even more so for *T. thermophila,* the number of transcripts was more steadily increasing with sequencing depth (Figure 4A, C). This indicates that more sequencing would not have rendered substantially more data for *P. tricornotum* and that the number of unique molecules in the final library was the limitation.

**Figure 4.**
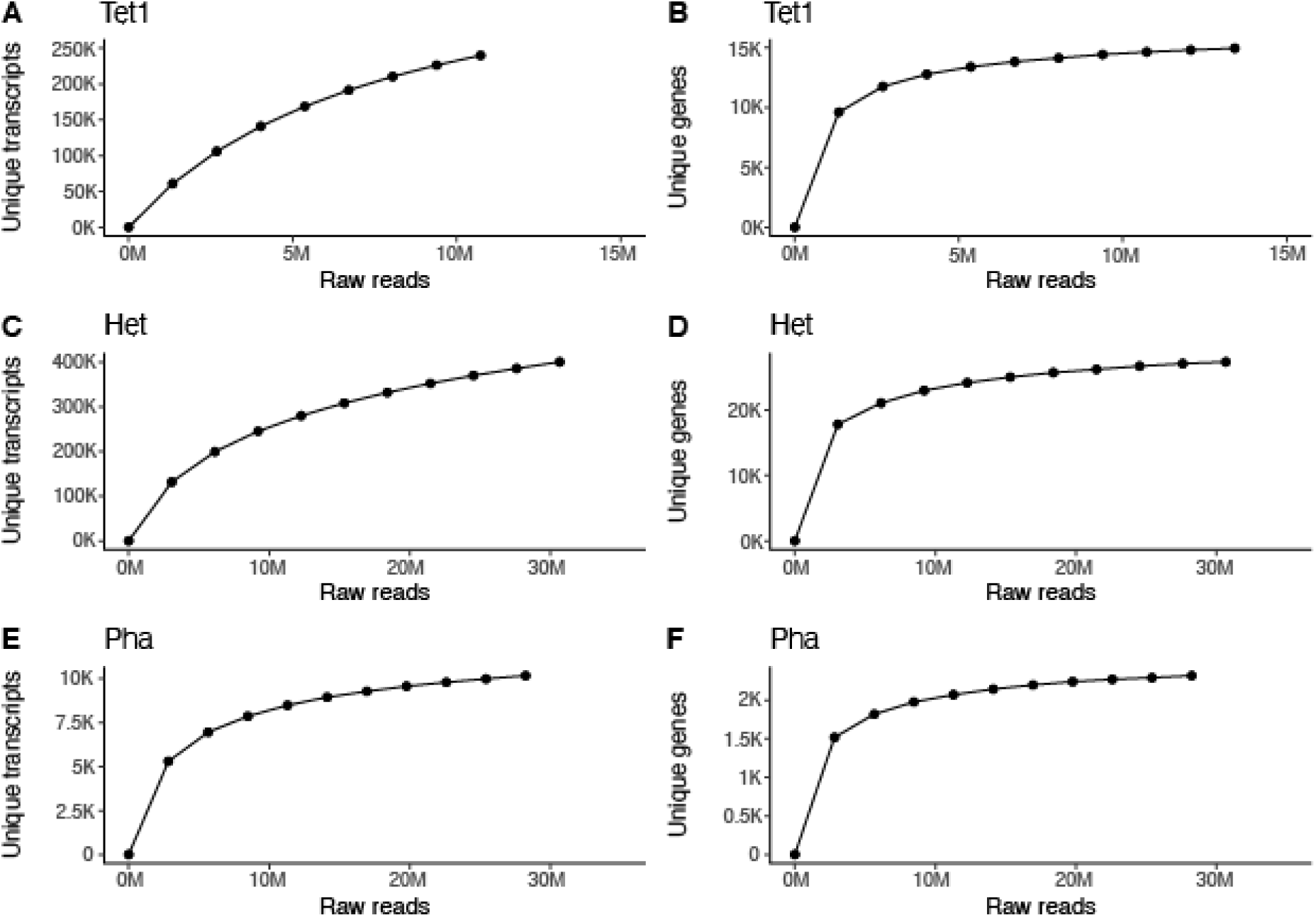
Saturation curves for each library *T. thermophila* (Tet1; A, B), *Heterocapsa* sp. (Het; C, D) and *P. tricornutum* (Pha, E, F) illustrating how many unique transcripts (A, C, E) and genes (B, D, F) that were detected as a function of sequencing depth. The sequencing depth (number of read pairs) are plotted on the x-axes and the unique transcripts or unique gene counts on the y-axes.

We investigated the ten most highly expressed genes, in spots that contained single cells, in more detail. In both the Tet1 and Tet2 libraries, genes for transmembrane proteins, papain family cysteine proteases and ribosomal proteins were the most highly expressed (Table 2). In the Het library, the most highly expressed genes encoded photosynthetic proteins peridinin-chl a protein, chloroplast proteins and major basic nuclear protein. We did not perform this analysis for the Pha library since gene counts in this case were very low.

**Table 2.** Top 10 most highly expressed genes (i.e. reference transcripts) in single-cell libraries of Heterocapsa sp. (Het) and T. thermophila (Tet1 and Tet2). Annotations were deduced by BLASTx searches against NCBI nr; reference transcript ID, protein description, organism and accession number of the best matches are given.

| Het<br>singles | reference transcript<br>ID | protein description | Organisms | gene<br>accession<br>ID |
| --- | --- | --- | --- | --- |
|  | TrINITY_DN4782_<br>c1_g1_i13 | peridinin-chl a protein | Heterocapsa pygmaea;<br>NCBI:txid35672 | CAC19481<br>.1 |
|  | TRINITY_DN4782_<br>_c1_g1_i7 | unnamed protein product | Heterocapsa pygmaea;<br>NCBI:txid35672 | CAE86223<br>56.1 |
|  | TRINITY_DN2086_<br>_c0_g1_i21 | major basic nuclear protein | Polarella glacialis;<br>NCBI:txid89957 | AAL61531<br>.1 |
|  | TRINITY_DN2086_<br>_c0_g1_i8 | major basic nuclear protein | Polarella glacialis;<br>NCBI:txid89957 | AAL61531<br>.1 |
|  | TRINITY_DN2086_c0_g1_i32 | major basic nuclear protein | Polarella glacialis;<br>NCBI:txid89957 | AAL61531.1 |
|  | TRINITY_DN2957_c0_g2_i4 | chloroplast ATPH isoform 5 | Heterocapsa triquetra;<br>NCBI:txid66468 | AAW80675.1 |
|  | TRINITY_DN4782_c1_g1_i4 | peridinin-chl a protein | Heterocapsa pygmaea;<br>NCBI:txid35672 | CAC19481.1 |
|  | TRINITY_DN2086_c0_g1_i18 | major basic nuclear protein | Pfiesteria piscicida;<br>NCBI:txid71001 | AAL61531.1 |
|  | TRINITY_DN2957_c0_g2_i3 | chloroplast ATPH isoform 5 | Heterocapsa triquetra;<br>NCBI:txid66468 | AAW80675.1 |
|  | TRINITY_DN2086_c0_g1_i14 | major basic nuclear protein | Pfiesteria piscicida;<br>NCBI:txid71001 | AAL61531.1 |
| Tet1<br>singles | reference transcript ID | protein description |  | gene<br>accession<br>ID |
|  | g13052.t1 | transmembrane protein,<br>putative | Tetrahymena thermophila SB210;<br>NCBI:txid312017 | XP_001032220.2 |
|  | g3965.t1 | papain family cysteine<br>protease | Tetrahymena thermophila SB210;<br>NCBI:txid312018 | XP_001027342.1 |
|  | g1442.t1 | hypothetical protein<br>TTHERM_001080487 | Tetrahymena thermophila SB210;<br>NCBI:txid312019 | XP_012655449.1 |
|  | g7083.t1 | transmembrane protein,<br>putative | Tetrahymena thermophila SB210;<br>NCBI:txid312020 | XP_012654551.1 |
|  | g3276.t1 | hypothetical protein<br>TTHERM_000760291 | Tetrahymena thermophila SB210;<br>NCBI:txid312021 | XM_012800270.1 |
|  | g25171.t1 | hypothetical protein<br>TTHERM_01125120 | Tetrahymena thermophila SB210;<br>NCBI:txid312022 | XP_001030140.1 |
|  | g16201.t1 | papain family cysteine<br>protease | Tetrahymena thermophila SB210;<br>NCBI:txid312023 | XP_012653036.1 |
|  | g10027.t1 | 60S ribosomal protein L31,<br>putative | Tetrahymena thermophila SB210;<br>NCBI:txid312024 | XP_001031042.2 |
|  | g162.t1 | ubiquitin-40S ribosomal<br>protein S27a | Tetrahymena thermophila SB210;<br>NCBI:txid312025 | XP_001025217.1 |
|  | g16902.t1 | 40S ribosomal protein S23 | Tetrahymena thermophila SB210;<br>NCBI:txid312026 | XP_001011429.2 |

| Tet2<br>singles | reference transcript<br>ID | protein description |  | gene<br>accession<br>ID |
| --- | --- | --- | --- | --- |
|  | g13052.t1 | transmembrane protein,<br>putative | Tetrahymena thermophila SB210;<br>NCBI:txid312022 | XP_00103<br>2220.2 |
|  | g3965.t1 | papain family cysteine<br>protease | Tetrahymena thermophila SB210;<br>NCBI:txid312023 | XP_00102<br>7342.1 |
|  | g1442.t1 | hypothetical protein<br>TTHERM_001080487 | Tetrahymena thermophila SB210;<br>NCBI:txid312024 | XP_01265<br>5449.1 |
|  | g7083.t1 | transmembrane protein,<br>putative | Tetrahymena thermophila SB210;<br>NCBI:txid312025 | XP_01265<br>4551.1 |
|  | g3276.t1 | hypothetical protein<br>TTHERM_000760291 | Tetrahymena thermophila SB210;<br>NCBI:txid312026 | XM_01280<br>0270.1 |
|  | g16902.t1 | 40S ribosomal protein S23 | Tetrahymena thermophila SB210;<br>NCBI:txid312027 | XP_00101<br>1429.2 |
|  | g14283.t1 | 60S ribosomal protein L36a | Tetrahymena thermophila SB210;<br>NCBI:txid312028 | XP_00101<br>0053.1 |
|  | g25171.t1 | hypothetical protein<br>TTHERM_01125120 | Tetrahymena thermophila SB210;<br>NCBI:txid312029 | XP_00103<br>0140.1 |
|  | g3569.t1 | Crystal Structure Of The<br>Eukaryotic 40s<br>Ribosomal Subunit In<br>Complex With Initiation<br>Factor 1. | Tetrahymena thermophila SB210;<br>NCBI:txid312030 | 2XZM_Z |
|  | g10027.t1 | 60S ribosomal protein L31,<br>putative | Tetrahymena thermophila SB210;<br>NCBI:txid312031 | XP_00103<br>1042.2 |

## Discussion

To the best of our knowledge, this study is the first to apply high-throughput parallel imaging and single-cell transcriptomics to microbial eukaryotic plankton. A major advantage of the MASC-seq method is the generation of images of the cells for which RNA is sequenced. This has great potential when studying uncultured microbial eukaryotes whose morphology and ecological roles are largely unknown. A microscope image of a cell can reveal information that cannot be interfered from genetic data alone, such as morphology, pigmentation, symbiotic relationships^29^, and by running the method on natural samples it has the potential to provide morphological data to microbial “dark matter”^8, 30^ for which only DNA sequence data exist to date.

Most single-cell transcriptomics protocols have been developed for and benchmarked on human and other mammalian cells but few have been tested on protists^16^. Planktonic protists are very diverse in terms of cell structure and thorough studies are needed to gain insights in a methods’ performance on different protist types. Therefore, our study represents a valuable contribution to the field.

The MASC-seq method worked fairly well on the ciliate (*T. thermophila*), and dinoflagellate (*Heterocapsa* sp.) species, with median values of 148 - 305 genes and 171 - 338 transcripts detected per cell for single-cell spots, and significantly more data for single-cell than background spots. Moreover, total expression levels displayed reasonable correlations between MASC-seq from cells and from bulk RNA. The diatom (*P. tricornutum*), however, gave much lower counts (5/8 genes/transcripts per cell) and expression for single-cell spots was not significantly different from the background spots. This is probably related to the diatom’s much smaller cell volume and lower number of mRNA molecules per cell^20^; the ratio between estimated number of mRNA molecules per cell and number of detected transcripts for single-cell spots was actually quite similar for the different species and libraries (156, 81, 271 and 183 for Pha, Het, Tet1 and Tet2, respectively). The reason why not more of the mRNA molecules of the cells were recovered is likely a combination of RNA degradation, incomplete diffusion out of the cells, binding to the underlying probes and cDNA synthesis, uneven amplification of the library, and insufficient sequencing depth. Which of these factors are more important is not clear, but our analysis indicated that sequencing depth was not the major limitation.

Although empty spots showed lower gene/transcript counts than spots with cells, the presence of background signal is still problematic since it indicates that a small amount of leakage between cells may occur, resulting in transcripts in some instances being assigned to the wrong cells. This issue will be particularly problematic when the method is run on microbial communities with species of varying sizes, since background signals from the larger cells will mask the foreground signal of the smaller. This can to some extent be alleviated by pre-selecting cells within a certain size range, for example by serial filtration. It is not clear to what extent the background signal comes from RNA already present in the suspension when the cells are smeared on the array, or are derived from diffusion from cells on the array after permeabilization with pepsin. The freeze-thaw cycle that was added before the cells were smeared could potentially contribute to RNA in the suspension, however we observe background signals also for the *Tetrahymena* libraries where this was not applied. Interestingly, the Tet2 library shows a much higher signal-to-background ratio than the Tet1 library, indicating that it is possible to further optimize conditions to lower the problem of background signal.

Another challenge of the protocol is obtaining an even distribution of cells over the array surface. Smearing of a cell suspension is a fast procedure but it can result in clumping of cells in different regions of the array. And even when cells are evenly distributed, only a fraction of the spots will harbor a single cell, and also these may have other cells directly outside in the surrounding array area. In the original MASC-seq publication^22^, an alternative approach of precision fluorescence-activated cell sorting (FACS) of cells into spots was also applied, which results in a larger fraction of single-cell spots and less background noise. However, this requires a specialized instrument and optimizations for different cell types, and most protists cannot be easily sorted by FACS. Another alternative to smearing could be to add the cells to the array in a larger suspension volume and performing a gentle swing bucket centrifugation of the slide as described in Colin et. al^31^. With this approach, cells would be distributed more evenly across the array and cell attachment may be improved and therefore loss during the different washing steps smaller. Furthermore, capture spot resolution and technologies continue to improve^32, 33^ reaching a resolution smaller than the diameter of a typical cell (0.5 - 1 µm), thereby increasing the likelihood that only one cell will overlap a spot.

In conclusion, our study shows that the MASC-seq approach has a potential for gaining parallel morphological and transcriptomic data for microbial eukaryotes in a high-throughput manner and hence may serve as a powerful tool for gaining novel insights into microbial dark matter.

## Methods

### Microarray slides

In this study two types of slides were used, optimization slides and library preparation slides; both slides were prepared on CodeLink-activated microscope glass slides (SurModics) as described by Ståhl et al.^34^. *Optimization slides* (Figure 1B) contained six array surfaces with uniform probes. The reverse transcription (RT) probes contained poly-T20VN oligonucleotides and were immobilized by printing on the array surface. Optimization slides are used for optimizing the key steps of the protocol: cell attachment, permeabilization, cell removal by analyzing the fluorescent cDNA ‘footprint’ that remained after successful RNA capture and removal of the cell (Figure 1A).

#### Library preparation slides

(Figure 1C) were printed in six identical 6.3 × 6.7 mm^2^ capture arrays. Each array includes 66 frame spots for orientation purpose and 1934 barcoded spots for capturing gene expression information. The spots have a diameter of ∼100 μm and are arranged in a centered rectangular lattice pattern: each spot has four surrounding spots with a center-to-center distance of 150 μm, forming a 310 μm x 325 μm rectangle^35, 36^. In each spot the capture probes share a barcode unique to that spot. During reverse transcription, the barcode is incorporated into the cDNA of all mRNAs captured on the spot (Figure 1D). This enables linking RNA-seq data to a spot on the array, and thereby the location of the cell from which the RNA originated. Each probe covalently immobilized on the glass surface has the following common 5′–3′ structure: 18-mer spot-unique positional barcode, 9-mer semi-randomized UMI^37^ and poly-20TVN capture region^38^. UMIs were utilized to identify unique transcripts per gene.

### Cell cultivation

We chose three common plankton types, a diatom (*P.tricornutum* CCAP 1055/1), a dinoflagellate (*Heterocapsa sp.* CCAP 1125/4) and a ciliate (*Tetrahymena thermophila* 630/1M). Both *P. tricornutum* and *Heterocapsa* were purchased from the Culture Collection of Algae and Protozoa (CCAP) from the Scottish Association for Marine Science (SAMS). Cells were grown and kept in L medium (*Heterocapsa sp.*), f/2 medium amended with silica (*P. tricornutum*) and PPY medium (*T. thermophila*). For each cell type we estimated growth curves based on microscopic counts (data not shown). Cells were harvested during the exponential phase and concentrated by centrifugation at 8000g for 5 min.

### RNA extraction from cell cultures

For each cell type we collected samples for bulk RNA. Cells were harvested on 5 µm polycarbonate filters (Whatman). Filters were placed in cryoprotectant tubes with beads (Zirconium oxide beads, Precellys), amended with 600 µl of RLT buffer (Qiagen) which contained 1% of 2-Mercaptoethanol, shock frozen in liquid nitrogen and stored at -80 °C until further processing. Prior to RNA extraction, samples were taken out of -80, thawed on ice and subjected to bead beating (1x 6000 rmp, Precellys Evolution homogenizer). *P. tricornutum* and *Heterocapsa* sp. samples were treated for 2 min and *T. thermophila* just vortexed vigorously. RNA from samples was further extracted with the RNeasy Qiagen kit (QIAGEN, Inc., Valencia, CA) following the kit instructions. RNA was re-eluted in 30 µl of nuclease free water.

### Optimization microarray experiments

We conducted a series of different tests on optimization microarray slides (Figure 1E). The basic workflow of optimization microarray experiment is illustrated in Figure 1A. The individual protocol steps that were optimized include: cell fixation, cell attachment, staining, permeabilization, cell removal and fluorescent cDNA footprint Subsequently, library preparation microarray slides were used.

#### Cell fixation

Two different fixatives were tested (formaldehyde and methanol) and their impact on fluorescent cDNA synthesis and final library quality evaluated (Supplementary figure 1). For this purpose, cells were chemically treated in three different fixatives during the exponential growth phase: fixation in formaldehyde, methanol and no fixation. For formaldehyde fixation, 2 ml of cell culture aseptically transferred into a microcentrifuge tube and supplemented with formaldehyde (36.5–38% (vol/vol); Sigma-Aldrich) to reach 2 % final concentration. Cells were incubated at room temperature for 10 min, centrifuged at 8000g for 5 min, the pellets were washed twice with 1xPBS and then shock frozen in liquid nitrogen. For methanol fixation, 1 ml of cell culture was aseptically transferred in a microcentrifuge tube and 1 ml of ice-cold methanol was added very slowly, in drops. Tubes were placed at -20 °C for a minimum of 30 minutes. Cells were washed in the same way as described for formaldehyde fixation. Additionally, unfixed cells were collected which were then fixated directly on the slide. Cells were washed twice with PBS as described above and pellets were shock frozen in liquid nitrogen (Figure 1A).

#### Freeze-thaw cycle

In order to improve permeabilization of *P. tricornutum* and *Heterocapsa sp.* a freeze-thaw cycle in liquid nitrogen and room temperature water was applied, which has previously been demonstrated to increase the RNA recovery^14^. Cells were collected, fixed and pelleted as described above. The pellet was first frozen in liquid nitrogen for 30 sec and dipped into room temperature water for 10 sec. The cycle was repeated 3 times and pellets stored at -80 until further processing. Freeze-thaw cycle was not used for *T. thermophila* since their cell membrane is not hard to permeabilized with enzymes.

#### Attachment, staining and imaging

Cells collected as described were placed on ice and diluted to a concentration of ∼1,000 cells μl^−1^ with cold 1xPBS. Before attaching the cells on the array, the whole slide was warmed at 37 °C for 1 min. Smearing was performed by taking 3 μl of the cell suspension and slowly pipetting the cells on the center of the array surface without touching the surface with the tip and maintaining a droplet by surface tension. Then, the cell solution was smeared with the side of a pipette tip (again careful not to touch the surface of the array) followed by attachment to the slide at 37 °C for 3 min (as detailed in^22^). Cells were afterwards stained with hematoxylin and eosin (H&E). Hematoxylin was added to cover cells attached to the slide (500 μl) and incubated for 4 min. Slide was washed by dipping in the beakers of nuclease-free water until the excess hematoxylin is washed away. Afterwards 500 μl of bluing buffer was added, incubated for 2 min and washed as described in the previous step. Finally, 500 μl of buffered eosine was added to the slide, incubated for 1 min and washed with nuclease free water. Slide that now contained stained cells was air dried and mounted with 1ml of 85% glycerol (Merck Millipore) and covered with a coverslip (Menzel-Glaser). Detailed staining protocol is described in Salménet al.^39^ Imaging was performed with Metafer Vslide scanning system (MetaSystems, Mannheim, Germany) installed on an Axio Imager Z2 LSM700 microscope (Carl Zeiss, Oberkochen, Germany). All images were taken with the 20 x Plan-Apochromat objective lens, and stitched using the VSlide software (v1.0.0). M^-2^. After imaging, the coverslip was removed from the glass slide by dipping in nuclease free water and subsequently in 80% Ethanol to remove glycerol.

#### Permeabilization, fluorescent cDNA synthesis and cell removal

Glass slides with arrays were put in a hybridization cassette (ArrayIT Corporation) with a rubber mask that separates each slide into six different subarrays to secure reaction chambers for each array and perform on-array reactions. Cells attached to each array were washed with 100 μl of 0.1 x saline-sodium citrate buffer (SCC) (Sigma-Aldrich) diluted with nuclease free water. Aliquots of 0.1% pepsin (Sigma-Aldrich) dissolved in 0.1 M HCl (Sigma-Aldrich) were pre-warmed at 37 °C for 5 min and then 70 μl of the solution was added into each array in order to permeabilize the cells. Cells of *P. tricornutum* and *Heterocapsa sp.* were permeabilized for 5 min and *T. thermophila* for 1 min. Pepsin was washed away with 100 μl of 0.1 x SCC buffer and 70 μl of reverse transcription (RT) mixture for fluorescent cDNA synthesis was added. The RT mixture contained: 1× First Strand Buffer (Invitrogen), 5 mM DTT (Invitrogen), 0.5 mM of each dNTP (Fisher Scientific) except dCTP which was 12.5 μM, and with additional inclusion of 25 μM Cy3-labeled dCTPs (PerkinElmer, Waltham, MA) in a 0.19 µg µl^−1^ BSA, 50 ng µl^−1^ actinomycin D (Sigma-Aldrich), 1% DMSO (Sigma-Aldrich), 20 U µl^−1^ Superscript III (Invitrogen) and 2 U µl^−1^ RNaseOUT (Invitrogen). The mixture was incubated overnight at 42 °C.

Each array was washed as described above and 70 μl of proteinase K (Qiagen) in PKD buffer (Qiagen) was added and the slide incubated at 56 °C for 1h and 30min with 300 rpm interval shake. After cell removal, slides were washed in 2xSCC which contained 0.1% SDS (Sigma Aldrich) at 50 °C for 10 min and subsequently washed with 0.2xSCC and 0.1xSCC for 1 min at room temperature. Arrays were mounted with 500 µl SlowFade Gold Antifade reagent (Invitrogen). Fluorescent footprints of the cell’s cDNA were imaged as described in the section “*Attachment, staining and imaging”*, only with fluorescent settings of the microscope (Cy3 filters). If the experiment was successful, fluorescence could be observed (Figure 1E). *Attachment test.* In order to test if the cells attach on the slide without falling off, additional tests were performed (Figure 1E). Cells were attached, stained, imaged, washed, permeabilized as described above and imaged again in order to inspect for movement or cell loss. Two images were compared and analyzed with imageJ^40^.

#### Permeabilization

Three different types of permeabilization methods and their outcome on fluorescent cDNA synthesis was tested, including pepsin^22^, lysozyme, and 0.01M HCL (Figure 1E, Supplementary table 1). By inspecting fluorescent footprint of cDNA, it was estimated that 5min of pepsin worked the best for *P. tricornutum* and *Heterocapsa sp.* while 1 min for *T. thermophila*.

#### Cell removal

Removing cells from the array of the slide is important in order to avoid false positive cDNA fluorescence footprint and contaminations during library preparation. To test cell removal and optimal concentration of proteinase K, a simple experiment was performed. Cells were attached, stained and imaged as described above and incubated with three concentrations of proteinase K in the PKD buffer (1:4, 1:5, and 1:6). The slides were imaged again to inspect if cell removal was successful (i.e., if the cells were not visible at the slide). Concentration of 1:4 and 1:5 successfully removed cells while 1:6 concentration did not remove all of the cells (results not shown).

### Library preparation

Once the protocol was optimized on optimization microarray slides, we used library preparation slides to produce libraries for sequencing. One library from each cell type (*P. tricornutum* (Pha), *Heterocapsa* sp. (Het)) and two libraries for *T. thermophila* (Tet1 and Tet2) were prepared for sequencing. Additionally, extracted RNA from each cell type was smeared on the slide array and prepared together with the cells. Total extracted RNA from corresponding cultures was used as a positive control to single cells and it was estimated how many µl of RNA we need to smear on the slide to have the amount of ∼ 2000 cells. During the optimization experiments methanol fixation provided the best outcome on library quality (Supplementary Figure 1), therefore this method was used for all subsequent experiments and library preparation.

#### Attachment, staining and imaging

Cells were attached, stained and imaged as described in the optimization microarray experiment.

#### Permeabilization, fluorescent cDNA synthesis and cell removal

Permeabilization mixture was prepared as described in the optimization microarray experiment. cDNA mixture for library preparation differed from optimization mixture in the way that it did not contain fluorescently labeled Cy3-dCTP. Mixture contained: 1× First Strand Buffer (Invitrogen), 5 mM DTT (Invitrogen), 0.5 mM of each dNTP (Fisher Scientific), 0.19 µg µl^−1^ BSA, 50 ng µl^−1^ actinomycin D (Sigma-Aldrich), 1% DMSO (Sigma-Aldrich), 20 U µl^−1^ Superscript III (Invitrogen) and 2 U µl^−1^ RNaseOUT (Invitrogen) and cDNA was synthesized over night at 42 °C. After the cDNA was synthesized, cells were removed with proteinase K in the PKD buffer (1:4 ratio) and subsequently washed as described in an optimization microarray experiment.

#### Probe release

Release of probes with mRNA–cDNA hybrids from the array surface was performed by adding 70 µl of release mix to the array chambers. Release mix was composed of 1× Second Strand Buffer (Invitrogen), 8.75 µM of each dNTP, 0.20 µg µl–1 BSA, and 0.1 U µl– 1 USER enzyme (NEB). Incubation was performed at 37 °C for 2 hours with interval shaking and 300 r.p.m. Release mixture was collected and transferred to a 96 well plate and placed at - 20 °C until further processing with an automatic robotic station.

#### Fluorescent imaging of spots

A fluorescence signal on library preparation arrays was obtained by adding 70 µl of reaction mixture containing 0.96× PBS, 0.2 µM Cy3-anti-A probe (Eurofins, [Cy3]AGATCGGAAGAGCGTCGTGT) and 0.2 µM Cy3-anti-frame probe (Eurofins, [Cy3]GGTACAGAAGCGCGATAGCAG). Incubation was performed at room temperature for 10 min. Arrays were washed with 2× SSC containing 0.1% SDS at 50 °C for 10 min, then with 0.2× SSC and 0.1× SSC at room temperature for 1 min, respectively. Subsequently, library preparation arrays were mounted with 500 µl of SlowFade Gold Antifade Reagent (Invitrogen) and covered with a coverslip. Fluorescence imaging, image stitching and extraction were performed following the procedure described in the section of optimization microarray experiment.

#### Image alignment

Image alignment of corresponding hematoxylin and eosin-stained cells and fluorescence images of spots were performed manually using Adobe Photoshop CC (Adobe).

### Automated library processing

Libraries were prepared with Magnatrix 8000+ (Nordiag), an eight-channel robotic workstation capable of running custom made scripts for in-tip magnetic bead separations^41^. Released mRNA–cDNA hybrids were transferred to the robotic workstation where they underwent second strand synthesis, end repair and *in vitro* transcription. Following transcription, sequencing adaptors are ligated to the RNA, and another round of cDNA synthesis is performed. Each step in the process is followed by a reaction clean-up step. Detailed descriptions of every reaction happening in the robotic station are present in^22, 39, 41^.

#### Final libraries amplification, indexing and sequencing

After clean-up, cDNA was amplified and indexed. The Optimum number of PCR cycles for final library amplification was estimated by qPCR in 10 µl of final volume. The qPCR reaction mixture contained 2 µl of purified sample, 1× KAPA HiFi HotStart Readymix (KAPA Biosystems), 0.5 µM PCR InPE1.0 primer (Eurofins, 5′-AATGATACGGCGACCACCGAGATCTACACTCTTTCCCTACACGACGCT CTTCCGATCT-3′), 0.01 µM PCR InPE2.0 primer (Eurofins, 5′-GTGACTGGAGTTCAGACGTGTGCTCTTCCGATCT-3′), 0.5 µM PCR Index primer (Eurofins, 5′-CAAGCAGAAGACGGCATACGAGATXXXXXXGTGACTGGAGTTC-3′), and 1× EVA green (Biotium). The qPCR (Bio-Rad CFX Real-Time system) programme was as follows: 1×, 98 °C 3 min; 25×, 98 °C 20 s, 60 °C 30 s, 72 °C 30 s. The PCR was performed in a reaction volume of 25 µl with the above programme and the established optimal number of cycles per sample. A final extension at 72 °C for 5 min was included. Indexed libraries were purified by an automated MBS robot^42^ and eluted in 20 µl of Elution Buffer (Qiagen). The library curves and average length were assessed using the DNA 1,000 kit (Agilent) on a 2,100 Bioanalyzer (Agilent), and the concentration was measured using the Qubit dsDNA HS Assay Kit (Life Technologies) following the manufacturer’s instructions. Indexed libraries were diluted to 2 nM and sequenced using the Illumina NextSeq platform applying paired-end sequencing. The forward read (R1) was sequenced for 31 bases, and the reverse (R2) for 121 bases.

### MASC-seq data analysis

Raw fastq files were processed through the ST pipeline^41^, which involves removal of duplicate reads, homopolymer stretches, and reads with low quality. For the purposes of this study, we modified the original code slightly, making it possible for the user to decide which attribute tag in the GFF file to use when counting reads. The modified code is available at https://github.com/NBISweden/st_pipeline/. The ST pipeline uses the transcript information present in the reverse reads (R2) and the barcode information, unique to a certain spot, present in the forward reads (R1) together with the UMIs to generate a matrix of counts where the genes are represented as columns and the spatial barcodes are represented as rows, and each matrix cell then represents the expression level of a given gene in a given spot. ST pipeline requires two files as an input: indexed genome or transcriptome of the sequenced species and corresponding GFF or GTF file. Genomes of *P. tricornutum*^43^ and *T. thermophila*^44^ were previously sequenced and downloaded with corresponding GFF files (https://mycocosm.jgi.doe.gov/Phatr2/Phatr2.home.html; http://ciliate.org/index.php/home/downloads). A reference genome for *Heterocapsa* sp. is not available so we performed a de-novo assembly from bulk RNA-seq to obtain the transcriptome as a reference (see below). In order to make the analysis as comparable as possible between the three species, we predicted transcriptomes for *P. tricornutum* and *T. thermophila* using their GFF files and the program gffread (v0.12.1^45^) with settings to exclude mRNAs that have no CDS features, and ran the ST pipeline in transcriptome mode for all three species. Ribosomal RNA (rRNA) sequences were identified in each reference transcriptome using barrnap (v0.9) with settings ‘--kingdom euk --reject 0.1’, and were subsequently removed. The ST pipeline computes the expression level of a gene (reference transcript) as the sum of R2 reads mapping to the gene, but in order to remove duplicates stemming from amplification of the same starting molecule, it sorts transcripts with the same UMI by strand and start position, then groups them if they are within 250 bp of each other, allowing 1 mismatch in the UMI.

Top 10 most highly abundant genes were extracted in R studio^53^ and BlastX was used to obtain gene annotations.

### Bulk RNA sequencing of Heterocapsa sp

RNA from *Heterocapsa* sp. Was extracted as described above and sent to SciLifeLab/NGI for library preparation and sequencing (Solna, Sweden). Libraries were prepared with Illumina TruSeq stranded mRNA kit with poly-A selection. Samples were sequenced on NovaSeq6000 with 151nt (Read1) -10nt (Index1)-10nt(Index2)-151nt(Read2) setup.

The bioinformatic analyses conducted on the bulk RNA-seq data were organized into a snakemake^46^ workflow available at https://github.com/NBISweden/LTS-A_Andersson_2010-MASCSEQ. The workflow uses snakemake version 6.0.5.

Adapters were trimmed using cutadapt (v3.2)^47^ using Illumina TruSeq adapters and minimum read length of 50. For sample P21817_104, the cutadapt maximum error rate was set to 0 in order to properly remove adapters. SortMERNA (v4.2.0)^48^ was used to filter reads into rRNA and non_rRNA partitions and was run with default settings and all rRNA databases. Read quality was assessed with fastqc (v0.11.9)^49^ and multiqc (v1.10)^50^.

Transcriptome assembly was performed with Trinity (v2.12.0)^51^ on the mRNA partition for each sample. Trinity was run in strand-specific mode (’--SS_lib_type FR’) and default settings. Coding regions were predicted on transcripts using TransDecoder (v5.5.0) with default settings and the ‘Universal’ genetic code. The resulting gff file from TransDecoder was then used to only keep assembled transcripts with a predicted coding sequence.

### Obtaining microarray spot coordinates for different groups of spots

In order to obtain the coordinates of the microarray spots, we used the program ST Spot Detector, that uses the bright field image of the array (that shows the hematoxylin and eosin-stained cells), in combination with a fluorescent image (where the array spots are highly visible since they contain Cy3 dye), for obtaining the array coordinates for each spot^52^. Spots with one cell (singles), two cells (doubles), >2 cells (clusters) and no cells (background) were manually selected. The output of the program is a file with the coordinates of selected spots that can be used together with a matrix of counts (output of the ST pipeline) to calculate gene expression coming from selected spots.

### Downstream data analysis

Downstream data analysis and generation of graphs were conducted in *R*^53^

## Author contributions

A.F.A. and R.A.F. conceived the study. R.A.F maintained the cultures, V.G. performed the experiments with support of S.S. and B.N. J.S. and B.S. conducted bioinformatics analyses. V.G and A.F.A. conducted downstream analyses. V.G and A.F.A. wrote the paper with contributions from all authors. M. L designed Figure 1. All authors read and approved the final version of the paper.

## Acknowledgements

This work was funded by Formas grant 2017-00694 to A.F.A and R.A.F; additional funds from Knut and Alice Wallenberg Foundation to R.A.F supplemented the project; R.A.F. acknowledges the technical support by Elina Viinamäki and Charlotta Ekblom for culture maintenance and Ryno Lawson for contributing to method development. B.S. and J.S are financially supported by the Knut and Alice Wallenberg Foundation as part of the National Bioinformatics Infrastructure Sweden at SciLifeLab. Computations and data handling were enabled by resources in project snic2020-15-29/snic2020-6-126 provided by the Swedish National Infrastructure for Computing (SNIC) at UPPMAX. The authors acknowledge support from the National Genomics Infrastructure (NGI) in Solna for assistance with transcriptomics sequencing. We thank Joakim Lundeberg and Sanja Vickovic for providing advice on spatial transcriptomics technology, Annelie Mollbrink for technical assistance, and Maliheh Mehrshad for providing comments that improved the manuscript.

## Data availability

Raw reads for MASC-seq together with output files of ST pipeline (matrix of counts) and spot files are available on Array Express and BioStudies with accession number E-MTAB-12261. Raw reads of Heterocapsa bulk RNA seq are available on Array Express and BioStudies with accession number: E-MTAB-12262. Images of cells corresponding to MASC-seq libraries are available on FigShare (https://figshare.com/s/650f1cff0036eca2d515) together with examples of optimization experiment (https://figshare.com/s/cff17c0cb8c88c11adba) and index transcriptome that was used as a reference for mapping of MASC-seq data (https://figshare.com/s/5621c7af9d57585ebe06).

