## Supplementary material for "Towards high-throughput parallel imaging and single-cell transcriptomics of microbial eukaryotic plankton"

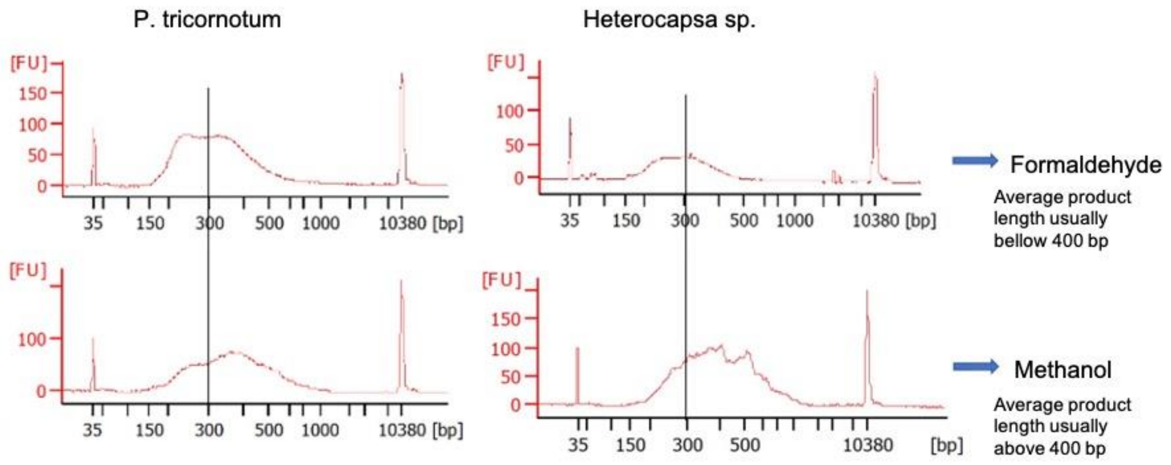


**Supplementary figure 1.** Bioanalyzer traces of final libraries showing product length when cells were fixed with formaldehyde and methanol. Upper and lower columns correspond to the same cells, collected at the same time points.

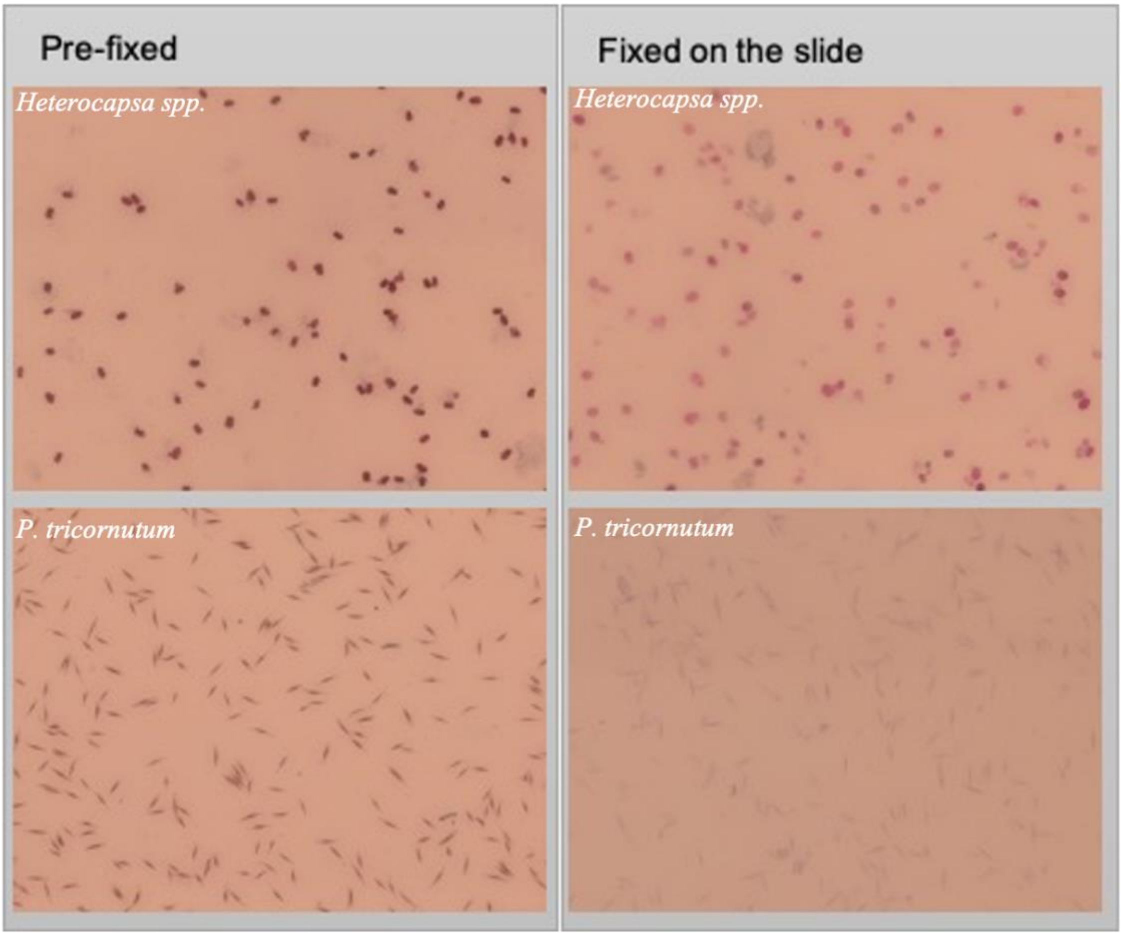

**Supplementary figure 2.** Cells of *Heterocapsa* sp. and *P. tricornutum* imaged after different ways of fixation.

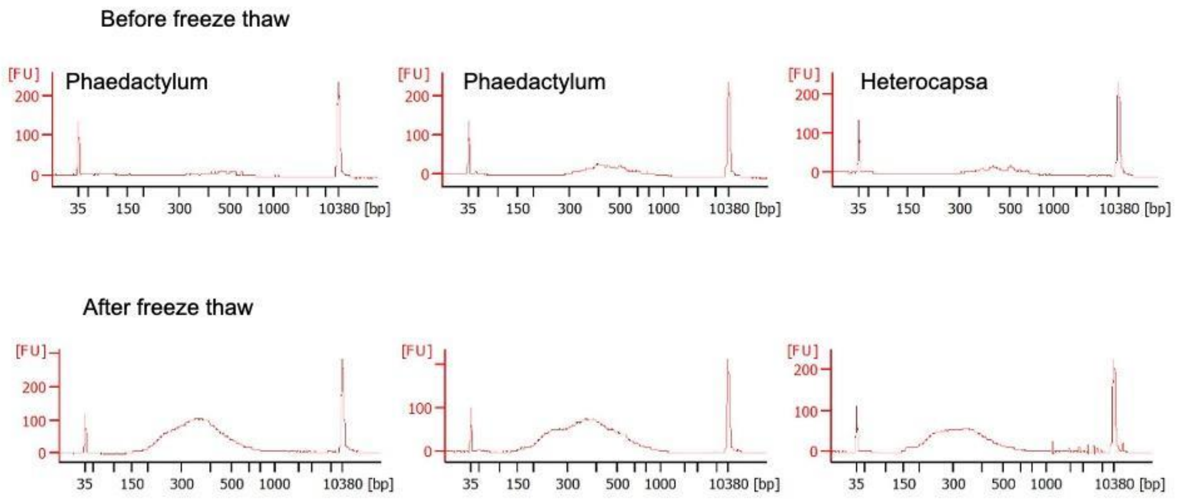


**Supplementary figure 3.** Bioanalyzer traces of final libraries before and after freeze-thaw cycling. Upper and lower columns correspond to the same cells, collected at the same time points.

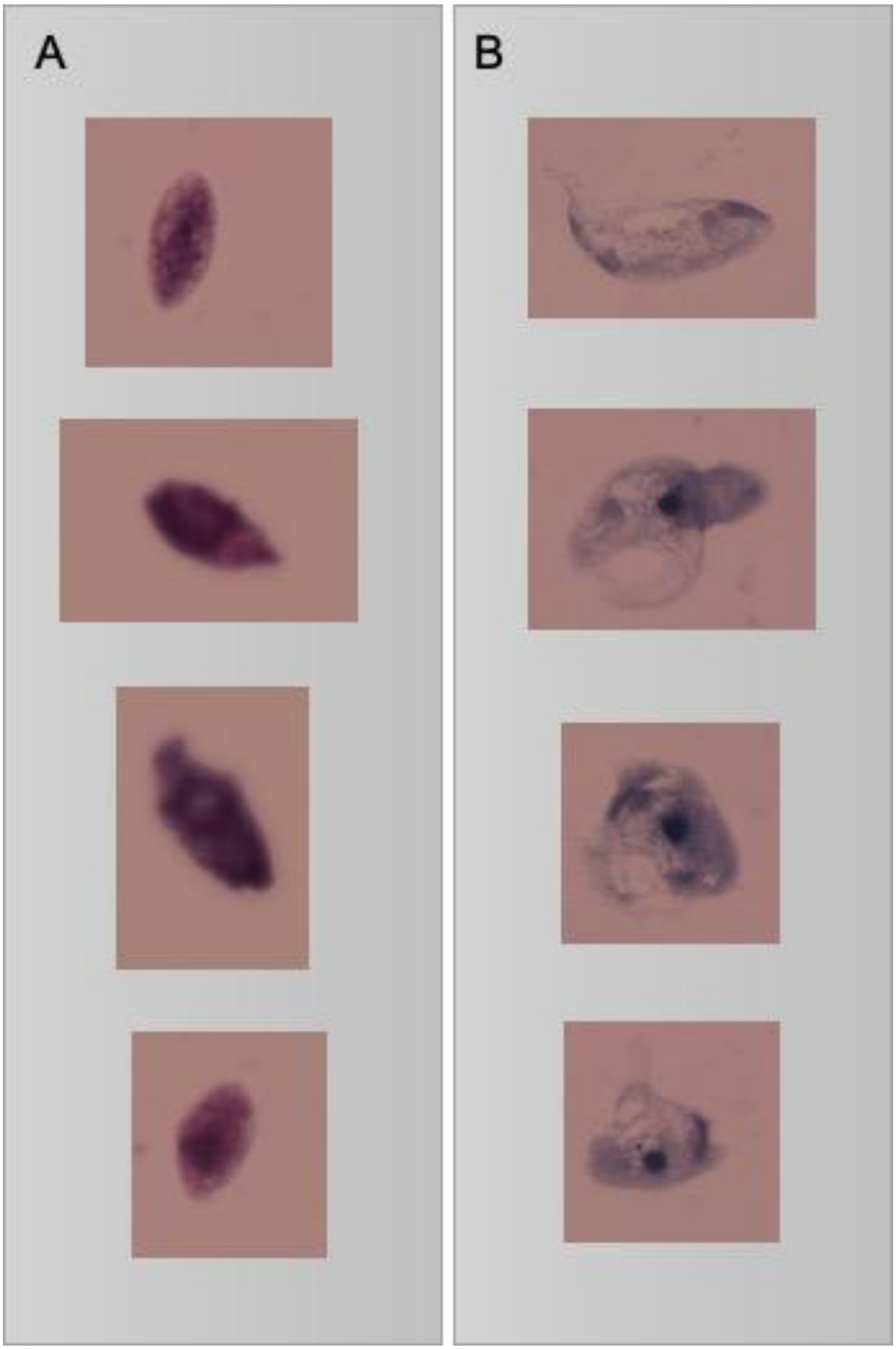


**Supplementary figure 4.** Examples of A) *T. thermophila* cells prefixed with methanol, attached to the slide and stained with hematoxylin and eosin; B) *T. thermophila* cells prefixed with methanol and then freeze-thawed in liquid nitrogen and room temperature water (3 times, attached on the slide and stained with hematotoxylin and eosin).

**Supplementary table 1.** Reagents and combinations used to test lysis on optimization slides

| **Reagent** | **Concentration** | **Temperature** | **Time** |
|---|---|---|---|
| Lysozyme solution* | 10 mg/ml | 37° C | 30 min<br>60 min |
| HCl | 0.01M | 37° C | 10 min<br>20 min |
| Pepsin# | 0.1 g $ml^{-1}$ | 37° C | 1 min<br>3 min<br>5 min<br>10 min |

* To prepare lysozyme solution add 100 mg lysozyme to 1 ml 0,5 M EDTA (pH 8), 1 ml 1M Tris/HCl (pH 8) and 8 ml nuclease free water
### Diluted 1:100 in 0.1M HCl
